# A systematic evaluation of data processing and problem formulation of CRISPR off-target site prediction

**DOI:** 10.1101/2021.09.30.462534

**Authors:** Ofir Yaish, Maor Asif, Yaron Orenstein

**Affiliations:** School of Electrical and Computer Engineering, Ben-Gurion University of the Negev, Beer Sheva, Israel

**Keywords:** CRISPR off-target, read count normalization, CHANGE-seq GUIDE-seq

## Abstract

CRISPR/Cas9 system is widely used in a broad range of gene-editing applications. While this gene-editing technique is quite accurate in the target region, there may be many unplanned off-target edited sites. Consequently, a plethora of computational methods have been developed to predict off-target cleavage sites given a guide RNA and a reference genome. However, these methods are based on small-scale datasets (only tens to hundreds of off-target sites) produced by experimental techniques to detect off-target sites with a low signal-to-noise ratio. Recently, CHANGE-seq, a new *in vitro* experimental technique to detect off-target sites, was used to produce a dataset of unprecedented scale and quality (more than 200,000 off-target sites over 110 guide RNAs). In addition, the same study included GUIDE-seq experiments for 58 of the guide RNAs to produce in vivo measurements of off-target sites. Here, we fill the gap in previous computational methods by utilizing these data to perform a systematic evaluation of data processing and formulation of the CRISPR off-target site prediction problem. Our evaluations show that data transformation as a pre-processing phase is critical prior to model training. Moreover, we demonstrate the improvement gained by adding potential inactive off-target sites to the training datasets. Furthermore, our results point to the importance of adding the number of mismatches between the guide RNA and the off-target site as a feature. Finally, we present predictive off-target in vivo models based on transfer learning from in vitro. Our conclusions will be instrumental to any future development of an off-target predictor based on high-throughput datasets.

## Introduction

Clustered regularly interspaced short palindromic repeats (CRISPR)/Cas9 system, originated in the immune defense system of archaea and bacteria [1, 2, 3, 4], is a groundbreaking tool for gene editing in a variety of cell types and organisms, and the preeminent gene-editing technology in recent years due to its low cost, high efficiency and effectiveness, and ease of use [5, 6, 7, 8, 9]. This system is composed of a Cas9 protein and an associated single guide RNA (sgRNA). The sgRNA guides the Cas9 to a target DNA sequence, and a double-strand break in the specific region is executed by the Cas9. The recognition of the DNA sequence is done via a complementarity of a 20-nucleotide (nt) sequence within the sgRNA to the genomic target upstream of a 3-nt protospacer adjacent motif (PAM) at its 3’ end. The CRISPR system is widely used in interrogating gene functions, and has wide applications in clinical detection, gene therapy, and agricultural improvement [10, 11, 12, 13, 14, 15].

Although the CRISPR/Cas9 system has many benefits, numerous studies have also shown that the mismatches between the designed sgRNA and DNA can be tolerated, resulting in cleavage of unplanned genomic sites, termed off-target sites. Therefore, detecting off-target sites in CRISPR gene editing is of high importance due to their disruptive effect, and designing sgRNAs with few off-target sites is desired [16, 17, 18]. Many experimental techniques were developed to detect off-target sites, including GUIDE-seq, Digenome-Seq, HTGTS, SITE-Seq, CIRCLE-Seq, and BLISS [19, 20, 21, 22, 23, 24]. However, those experimental techniques are expensive to apply, time-consuming, and produce noisy measurements that result in many false negatives.

To overcome these limitations, various computational methods have been developed to predict off-target sites. These methods can be split into three categories: alignment-based, ruled-based, and data-driven-based. Many data-driven methods are based on machine learning, such as CRISTA, DeepCrispr, Elevation, among many others [25, 26, 27, 28, 29, 30, 31, 32]. Still, all these methods were trained using relatively small experimental data, i.e. less than 10 sgRNAs per dataset with only tens of active off-target sites in total [33], resulting in inaccurate models for off-target prediction. For example, in a recent study, a state-of-the-art ensemble model was trained on a benchmark dataset and tested in cross-validation [34]. We gauged the model’s performance, and it achieved an average area under the precision-recall curve (AUPR) over 16 sgRNA targets of only 0.31 ± 0.2. In addition, most methods solve an off-target classification problem, with only few methods aiming to predict off-target cleavage efficiency. Moreover, most studies evaluate prediction performance on all sgRNAs together, while most applications use each sgRNA separately, and hence such evaluations are not informative. Solutions to the questions of how to process off-target sites data, and how to formulate the off-target site prediction problem, were limited due to the quality and size of available datasets.

Recently, a new dataset of unprecedented scale and accuracy was produced for both off-target sites *in vitro* and *in vivo* [35]. This dataset contains 202,044 *in vitro* off-target sites across 110 sgRNAs, and 1,702 *in vivo* off-target sites across 58 sgRNAs, which are a subset of the 110 sgRNAs. This abundance of data introduced a unique opportunity to systematically evaluate the problem definition and data handling of off-target site prediction. Such evaluations will be critical for any future development of accurate off-target predictors.

To take advantage of this opportunity, we exploit the new dataset to develop different model variants to predict off-target sites and to predict off-target cleavage efficiency and to perform an extensive comparison between the models. We demonstrate how our predictive models trained on in vitro data are able to make predictions on in vivo data more accurately than the in vitro measurements themselves. Furthermore, we provide a proof-of-concept that a regression model trained with inactive off-target sites, which were not obtained in the experimental dataset, improves the prediction over training only on off-target sites obtained in the experiment. In addition, we show the necessity of data transformation as a data pre-processing stage before training the regression models. Moreover, we approximate a nontrivial function between the number of sgRNA-DNA mismatches, termed *distance,* and the off-target cleavage efficiency. We also point out the distance feature’s importance for predicting both the off-target sites and off-target cleavage efficiency. Finally, we are the first, as far as we know, to present predictive off-target *in vivo* models based on transfer learning from *in vitro* data.

## Materials and Methods

### Dataset and prepossessing

#### CHANGE-seq and GUIDE-seq datasets

We used two high-throughput CRISPR off-target datasets, one *in vitro* and one *in vivo* (Figure 1a). CHANGE-seq is a scalable, highly sensitive, and unbiased assay for measuring the genome-wide activity of CRISPR/Cas9 nucleases *in vitro* [35]. The developers of CHANGE-seq applied their method to 110 sgRNA targets across human primary T cells and identified 202,044 off-target sites with up to six mismatches compared to the sgRNA. GUIDE-seq is an *in vivo*, low-throughput, and unbiased approach for global detection of DNA double-stranded breaks introduced by CRISPR RNA-guided-nucleases [19]. The efficiency of GUIDE-seq is effected by chromatin accessibility and other epigenetic factors [36]. The developers of CHANGE-seq conducted GUIDE-seq experiments to compare their measurements with the measurements produced by the CHANGE-seq assay. They applied GUIDE-seq to 58 sgRNAs, which were also tested in the CHANGE-seq experiment, and identified 1,702 off-target sites (Figure 1b). In both CHANGE-seq and GUIDE-seq datasets, the off-target cleavage efficiency is quantified as the number of read counts. All of the models in this study were trained on the CHANGE-seq dataset only. For the models produced by transfer learning, additional training was performed on GUIDE-seq experimental data.

**Figure 1:**
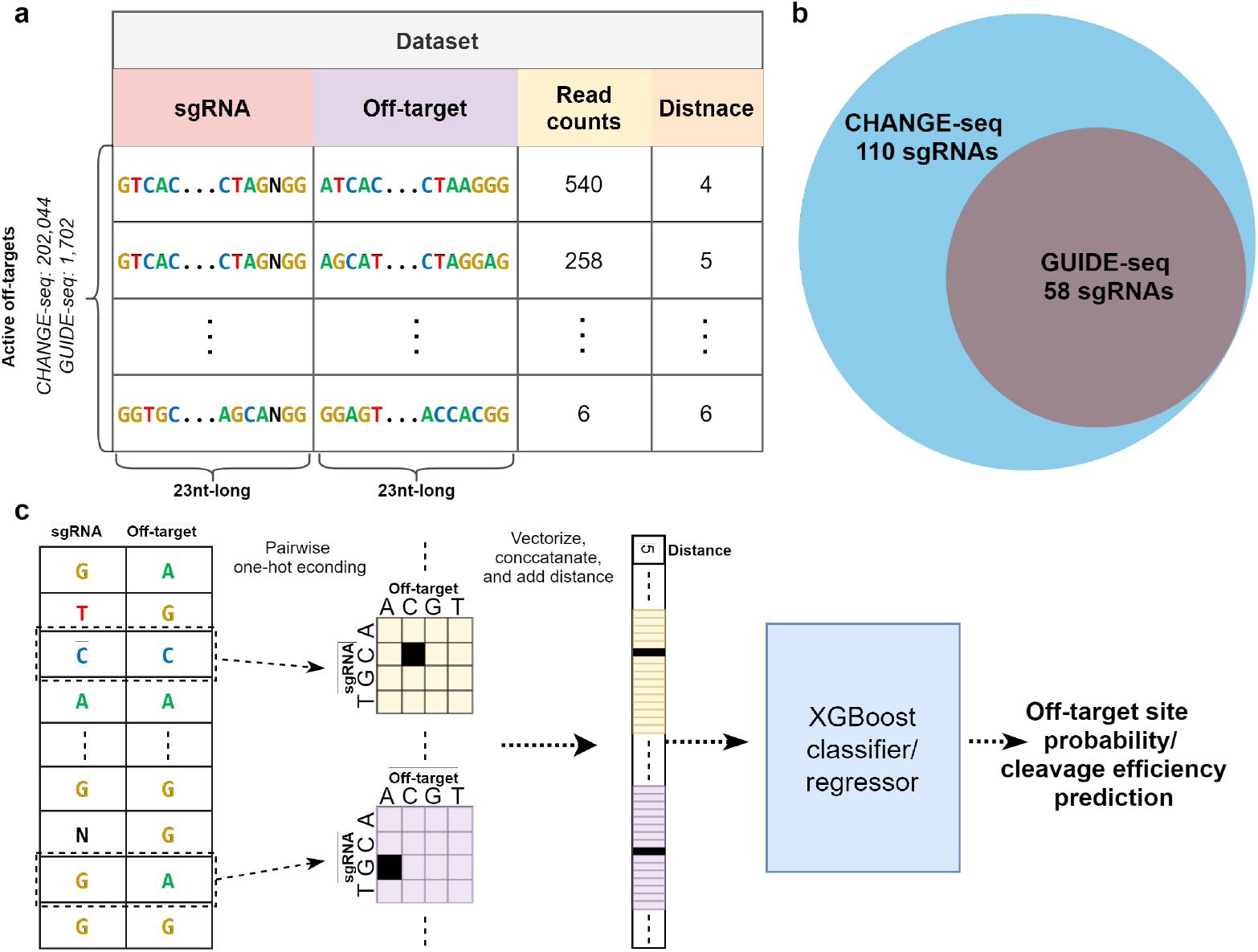
Data and computational model overview. **(a-b)** Illustration of the CHANGE-seq and GUIDE-seq datasets. **(a)** The developers of CHANGE-seq tested the off-target sites of 110 sgRNAs across human primary T cells. The CHANGE-seq dataset contains, across 110 sgRNAs, 202,044 active off-target sites with up to six mismatches compared to the sgRNA. In addition, they tested the off-target sites of 58 of the 110 sgRNAs by GUIDE-seq. The GUIDE-seq dataset contains, across 58 sgRNAs, 1,702 active off-target sites with up to six mismatches compared to the sgRNA. For each off-target site the number of read counts and the number of mismatches, termed distance, are reported. **(b)** The off-target sites of 58 out the 110 CHANGE-seq sgRNAs were also tested in a GUIDE-seq experiment. **(c)** Overview of the XGBoost predictive model. Given a pair of sgRNA and off-target sequences, the sequences are being processed using pairwise one-hot encoding. Then, a concatenation of the flattened pairwise one-hot encoded matrices and the distance feature (if required) is provided as input to the XGBoost classifier/regressor to predict the off-target site probability/cleavage efficiency.

#### Generating the active and inactive off-target sets

To train and test our classification and regressions machine-learning models, we defined an active off-target sites set (i.e., positive set) and an inactive off-target sites set (i.e., negative set). On CHANGE-seq, as done in the original study, the active off-target sites are the experimental off-target sites cleaved *in vitro* with read counts greater than 100. The remaining CHANGE-seq samples (i.e., with read counts less than 100) were considered to be sampling noise and were excluded from all sets. The CHANGE-seq dataset contains off-target sites with DNA/RNA bulges, which we filtered out to simplify the computational model. On GUIDE-seq, we considered as active off-target sites all experimentally obtained off-target sites. To generate the inactive off-target sites set, we used Cas-OFFinder [37].Given a sgRNA and a reference genome, Cas-OFFinder detects all potential off-target sites of Cas9 RNA-guided endonucleases with up to six mismatches compared to the sgRNA. From the potential off-target sites obtained by Cas-OFFinder, we filtered out all experimentally identified sites. Overall, for the CHANGE-seq dataset, we generated active and inactive sets of sizes 67,476 and 2,806,152, respectively, over 110 sgRNAs. For the GUIDE-seq dataset, we generated active and inactive sets of sizes 1,702 and 1,476,301, respectively, over 58 sgRNAs.

### Machine-learning models

#### Sequence encoding

In our machine-learning models, off-target activity is predicted based on the sgRNA and off-target sequences, or the number of mismatches between them, termed distance, or both. We encoded the sgRNA and off-target sequences in the same way as in the CHANGE-seq study. Given a pair of sgRNA and off-target sequences, *S_t_* and *S_o_,* respectively, the pair of nucleotides in position *i* of the sequences is encoded into a 4 × 4 matrix as follows:

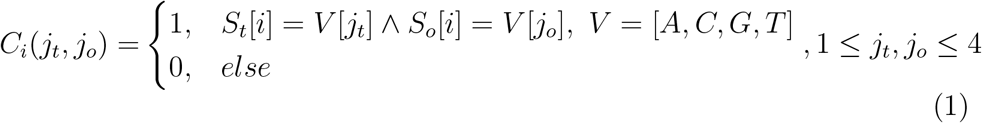

To create the sequence feature vector of one pair of sequences, each matrix representing one pair of nucleotides is flattened, and all matrices are concatenated into a single vector in the order of the nucleotides in the sequences. This results in a feature vector of size 368. The distance feature may be concatenated into the sequence feature vector, depending on the model we tested.

#### Classification and regression models

We developed two types of predictive models. The first model is a classification model to predict the probability of an off-target site to be cleaved (i.e., the classification task). The second model is a regression model to predict the off-target cleavage efficiency of an off-target site (i.e., the regression task). We implemented multiple variations of the models: models based only on the distance feature (i.e., <classification/regression>-dist), models based only on the sequence features (i.e., <classification/regression>-seq), and models based on both distance and sequence features (i.e., <classification/regression>-seq-dist).

For the regression models, we tested training on *log*(*x* + 1) data transformation where *x* is the read count. We considered using Box-cox transformation as performed in CRISPR-Net and Elevation studies [28, 30], or Yeo–Johnson transformation; however, we found them infeasible to apply due to our evaluation approach, which assumes no connection between the training and the test sets, whereas in previous studies, the data transformation, which learns its parameters from the data, was applied on all the data. In addition, as done in previous studies [28, 30, 38, 27, 26], we tested the option of adding the negative set to the training set of the regression model with label zero.

Our proposed predictive models are based on gradient boosting, a machinelearning method, which can solve regression and classification tasks (Figure 1c). It produces a predictive model in the form of an ensemble of weak decision trees [39]. We used XGBoost Python library in our implementation [40].

#### Model training

From the CHANGE-seq dataset, as mentioned above, we derived 67,476 positives samples and 2,806,152 negatives samples, which is an extremely imbalanced dataset. Training machine-learning models on an imbalanced dataset is usually inefficient [41, 42, 43]. Therefore, we used the XGBoost sample weight option to handle the data imbalance problem for both the classification and regression models. For each set, we defined its weight to be proportional to the size of the other set. For example, the weight of the active set is set to

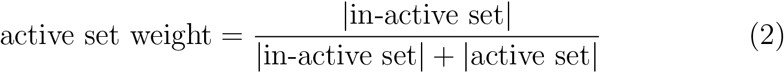

To train our XGBoost model, we used 1000 tree estimators with a max depth of 10 and a learning rate of 0.1.

#### Model evaluation

To assess how the predictions of our predictive models will generalize on an independent dataset, we used cross-validation for training and testing the models. We trained models using CHANGE-seq in an leave-11-sgRNAs-out cross-validation technique. This technique randomly partitions the CHANGE-seq dataset of 110 sgRNAs into 10 non-sgRNA-overlapping sets, i.e., each set contains samples of different 11 sgRNAs. In each iteration, of the 10 sets, a single set is retained as the test set, and the remaining 9 sets are used as the training data. To evaluate the performance on the GUIDE-seq dataset, for each sgRNA we used the model that was trained on all sets, excluding the set containing this sgRNA.

To evaluate the performance of the models on the classification task, we calculated AUPR, which is typically used to measure classification performance on an imbalanced dataset. To evaluate the performance of the models on the regression task, we used the Pearson correlation on the positive set read counts. In both evaluations, we measured the performance on each sgRNA separately.

#### Transfer learning from CHANGE-seq to GUIDE-seq

We suggest two approaches to train a model based on transfer learning. The first approach uses a pre-trained model on CHANGE-seq, and then continues the training on the GUIDE-seq dataset by adding new trees to the XGBoost model, which we denote *GS-TL-add*. The second approach updates the pre-trained trees to fit the GUIDE-seq dataset. We denote this approach *GS-TL-update*. We compared the two approaches to a model trained on the GUIDE-seq dataset without using a pre-trained model, which we denote *GS*. We trained the pre-trained models on the 52 CHANGE-seq sgRNAs that were not tested in a GUIDE-seq experiment, and left the 58 GUIDE-seq sgRNAs for the transfer learning and testing the obtained models.This was done to evaluate the ability of a pre-trained model to fit to GUIDE-seq experimental data, not containing the same sgRNAs.

In addition, to evaluate the number of GUIDE-seq experiments needed to train a model in a transfer learning approach, we randomly picked 10 experiments out of the 58 GUIDE-seq sgRNAs for testing the models. The rest of the GUIDE-seq sgRNAs were used for training the models in the transfer learning. For each number of GUIDE-seq experiments, 1 ≤ *i* ≤ 48, we randomly pick a set of *i* experiments and perform the transfer learning on them. We repeated this process 10 times for each such number for robust evaluation.

## Results

### Read counts log transformation improves prediction performance

We first examined CHANGE-seq read counts distribution (Figure 2a). The read counts distribution is highly skewed, i.e., there are many low read counts, and only a few read counts in the thousands. Therefore, we tested the effect of applying log transformation on the read counts. This transformation resulted in a much more balanced distribution (Figure 2b). To test whether this data transformation enhances the performance of the predictive models, we trained regression models on the training data with and without the log transformation, and tested their performance on both the classification and regression tasks.

**Figure 2:**
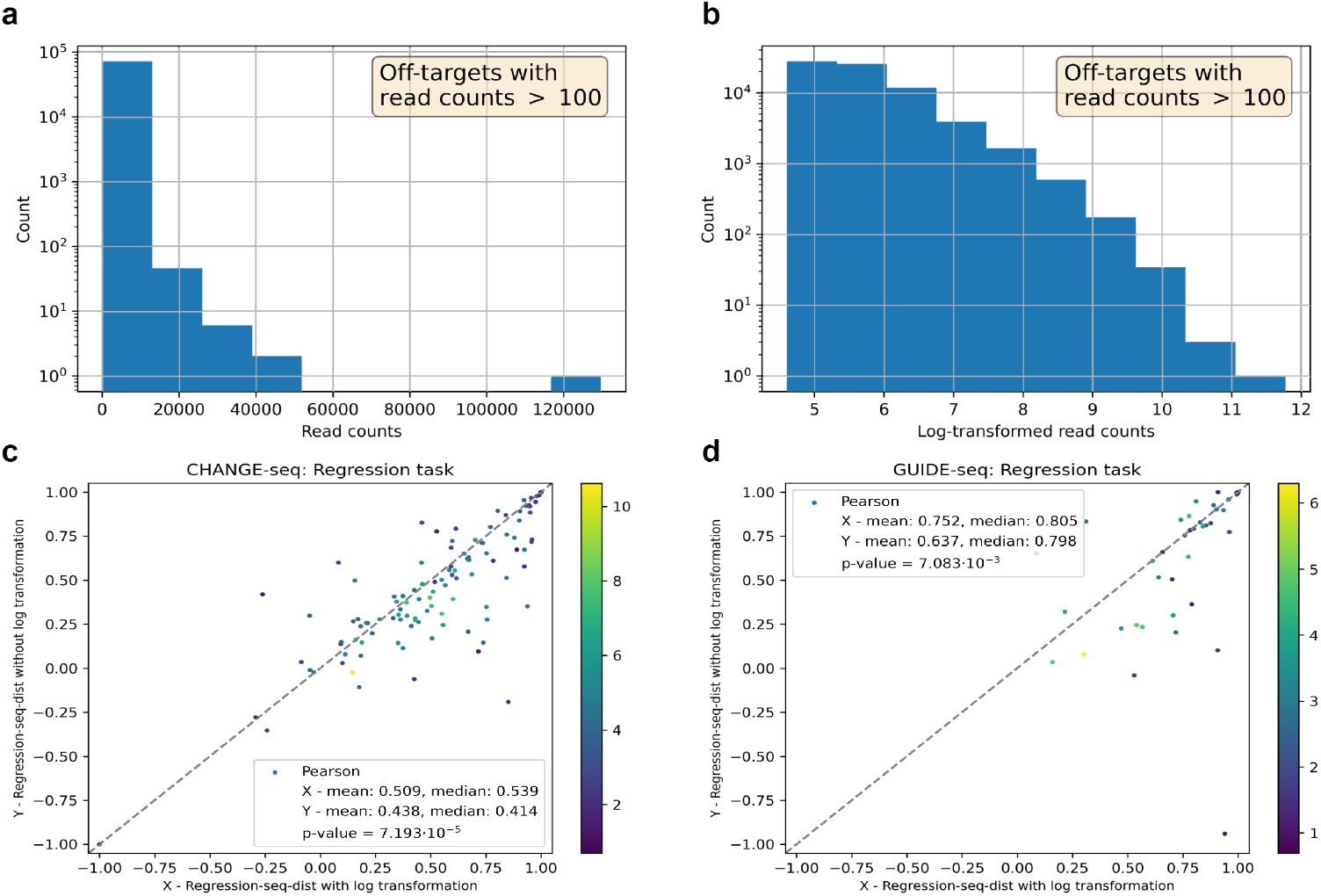
Read counts log transformation evaluation. **(a-b)** CHANGE-seq read counts distributions filtered to off-target sites with read counts greater than 100. **(a)** Read counts distribution. **(b)** Log-transformed read counts distribution. **(c-d)** Comparison of the regression-seq-dist model trained with log transformation and without. Using a leave-11-sgRNAs-out cross-validation, we evaluated the performance for the regression task on each sgRNA individually. Performance evaluation is preformed on the CHANGE-seq dataset (c), and the GUIDE-seq dataset (d). Performance was gauged by Pearson correlation between measured and predicted read counts. Each point in the scatter plots represents performance evaluated on a sgRNA. The color of each point in the scatter plots represents the log of the number of active off-target sites experimentally detected for this sgRNA. The reported p-value was computed by Wilcoxon signed-rank test.

We performed a regression performance comparison of the models on the CHANGE-seq and GUIDE-seq datasets (Figures 2c,d). On the CHANGE-seq dataset, the regression-seq-dist model with log transformation achieved the best average Pearson correlation of 0.509 compared to 0.438 of the same model without log transformation (p-value= 7.19 · 10^-5^). On the GUIDE-seq dataset, the regression-seq-dist model with log transformation achieved an average Pearson correlation of 0.752 compared to 0.637 of the same model without log transformation (p-value= 0.007). The same trend was observed when evaluating the models in the classification task (Supplementary Figures S1a,b).

Overall, we observed that the performance of the regression models trained on data with log transformation of the read counts is much improved compared to the performance of models trained on the original read counts. Therefore, we trained the regression models with log transformation on the read counts in all following comparisons. The full results, on each sgRNA, are reported in Supplementary Tables S1 and S2.

### Including potential off-target sites with no reads in regression model training improves prediction performance

When training a regression model to predict off-target cleavage efficiency, we can either train on only active off-target sites, which were experimentally detected in a CHANGE-seq experiment [35], or train on a set that includes both active and inactive off-target sites, which were obtained using Cas-OFFinder [37] and assigned a zero read count label. To assess the two options, we trained two regression models: one including and the other excluding the inactive set. We then tested their performance in both classification and regression tasks.

We performed a regression performance comparison of the models on CHANGE-seq and GUIDE-seq datasets (Figures 3a,b). Note that the evaluation is done only on the active set. On the CHANGE-seq dataset, the regression-seq-dist model trained with the inactive set achieved the best average Pearson correlation of 0.509 compared to 0.439 of the same model trained without the inactive set (p-value= 4.70 · 10^-4^). On the GUIDE-seq dataset, the regression-seq-dist model trained with the inactive set achieved an average Pearson correlation of 0.752 compared to 0.614 of the same model trained without the inactive set (p-value= 1.99 · 10^-4^). The same trend was observed when evaluating the classification performance of the models (Supplementary Figures S2a,b).

**Figure 3:**
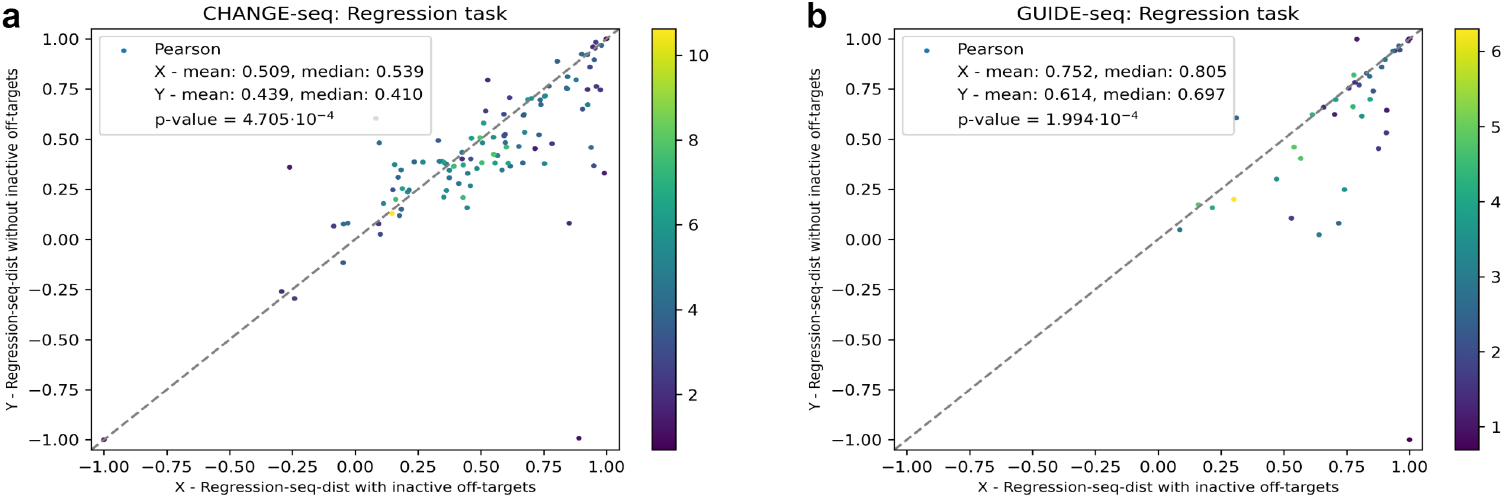
Training on both inactive and active off-target sites evaluation. **(a-b)** Comparison of the regression-seq-dist model trained only on active off-target sites and trained on both active and inactive off-target sites. Using a leave-11-sgRNAs-out cross-validation, we evaluated the performance for the regression task on each sgRNA individually. Performance evaluation is preformed on the CHANGE-seq dataset (a), and the GUIDE-seq dataset (b). Performance was gauged by Pearson correlation between measured and predicted read counts. Each point in the scatter plots represents performance evaluated on a sgRNA.The color of each point in the scatter plots represents the log of the number of active off-target experimentally detected for this sgRNA. The reported p-value was computed by Wilcoxon signed-rank test.

In summary, training a regression model on both the active and inactive sets improves performance in both classification and regression tasks. Therefore, we used the regression models trained with the inactive set in all following comparisons. The full results, on each sgRNA, are reported in Supplementary Tables S1 and S3.

### The combination of sequence and distance features achieves best prediction performance

Next, we aimed to determine which model performs best in predicting active off-target sites, and which one performs best in predicting off-target cleave efficiency. For this goal, we evaluated the performance of the classification models trained on sequence features with and without the distance feature, and of the equivalent regression models on both CHANGE-seq and GUIDE-seq datasets.

First, we compared the classification performance of the models (Figures 4a,b). On the CHANGE-seq dataset, the classification-seq-dist model achieved the highest average AUPR of 0.643 compared to 0.551, 0.475, 0.319 of regression-seq-dist, classification-seq, and regression-seq models, respectively (p-values < 7.66 · 10^-15^). On the GUIDE-seq dataset, the regression-seq-dist model achieved an average AUPR of 0.658. The regression-seq-dist model outperformed the classification-seq and regression-seq models, which achieved an average AUPR of 0.584 and 0.540, respectively (p-values < 1.94 · 10^-5^). The classification-seq-dist model obtained results on par with the regression-seq-dist and classification-seq models with an average AUPR of 0.605 (p-values > 0.1).

**Figure 4:**
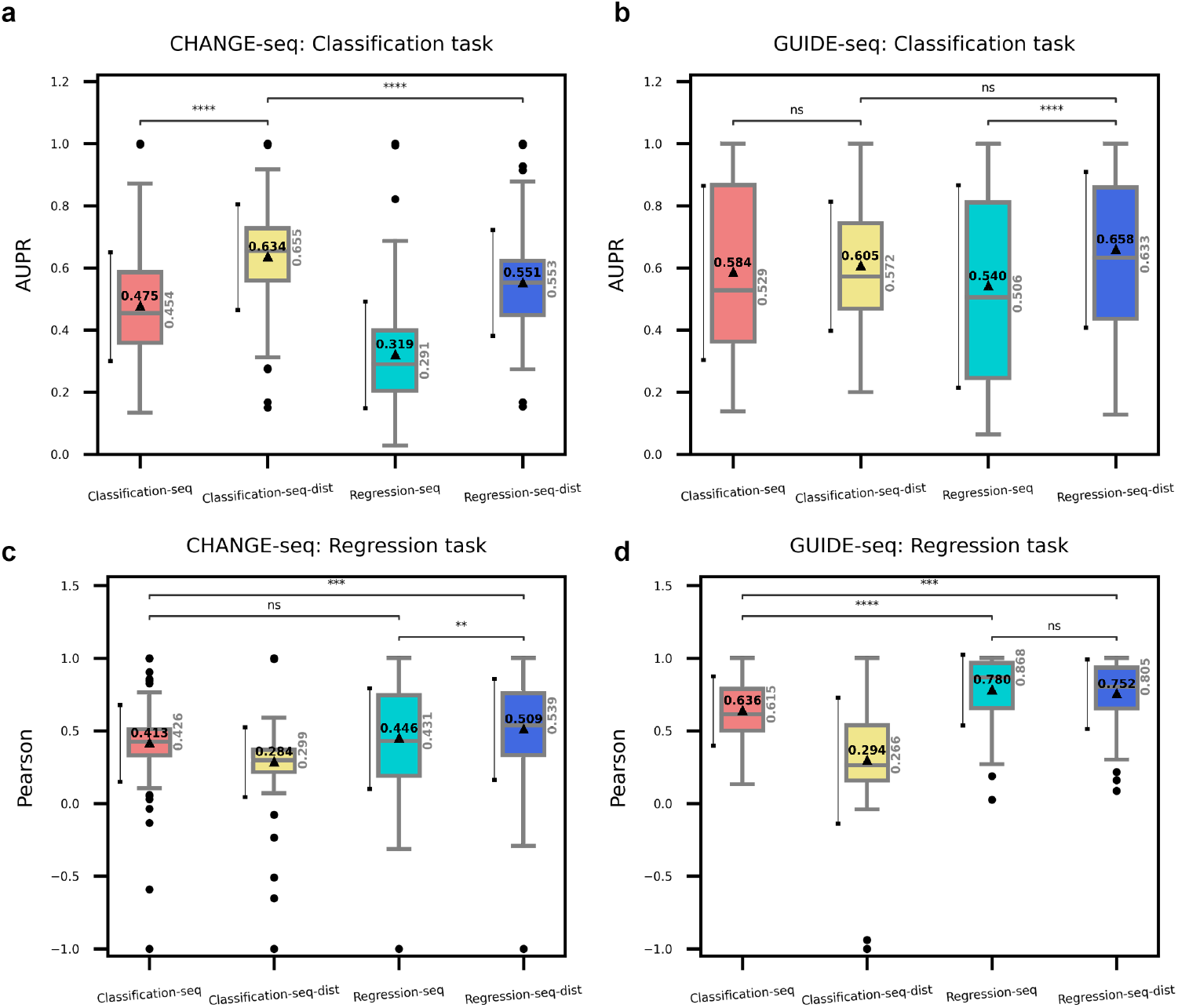
Classification and regression models comparison. **(a-d)** Comparison of <classification/regression>-seq, and <classification/regression>-seq-dist models in the classification and regression tasks. Performance was gauged by AUPR for classifying active off-target sites and Pearson correlation of measured and predicted off-target read counts. The average AUPR and Pearson correlation are denoted by a triangle within each box plot. Error bar of one standard deviation is located on the left of each box plot. The median AUPR and Pearson correlation values are reported in grey on the right of each box plot. Statistical significance via Wilcoxon signed-rank test denoted by: ns 5 · 10^-2^ < *p* ≤ 1; ** 1 · 10^-3^ < p ≤ 1 · 10^-2^; *** 1 · 10^-4^ < p ≤ 1 · 10^-3^; **** p ≤ 1 · 10^-4^. **(a-b)** Performance evaluation of the classification task on the CHANGE-seq dataset (a), and the GUIDE-seq dataset (b). **(c-d)** Performance evaluation of the regression task on the CHANGE-seq dataset (c), and the GUIDE-seq dataset (d).

Second, we compared performance of the models in predicting off-target cleavage efficiency (Figures 4c,d). On the CHANGE-seq dataset, the regression-seq-dist model achieved the highest average Pearson correlation of 0.509 compared to 0.446, 0.413, 0.284 of regression-seq, classification-seq, classification-seq-dist models, respectively (p-values < 4.87 · 10^-4^). On the GUIDE-seq dataset, the regression-seq model achieved an average Pearson correlation of 0.780. The regression-seq model outperformed the classification-seq and classification-seq-dist models, which achieved an average Pearson correlation of 0.636 and 0.294, respectively (p-values < 2.32 · 10^-5^). The regression-seq-dist model obtained results on par with the regression-seq with an average Pearson correlation of 0.752 (p-value=0.965).

Overall, on both CHANGE-seq and GUIDE-seq, we observed that for the classification task, the best model is the classification-seq-dist model, and for the off-target cleavage efficiency prediction task, the best model is the regression-seq-dist model. Surprisingly, in the classification task the regression-seq-dist model achieved high performance, particularly on the GUIDE-seq dataset. Furthermore, the classification-seq model obtained much higher performance in the regression task than the classification-seq-dist model. However, in most cases, the results point to the importance of the distance feature, and to the preferred usage of combined sequence and distance features. The full results, on each sgRNA, are reported in Supplementary Tables S1 and S4.

### Evaluating regression task performance of models trained with the distance feature only

Following our results pointing to the importance of the distance feature, we set out to test the prediction performance of models with only the distance feature. To study the impact of the distance feature, we defined the regression-dist model, which is trained only with the distance feature, and compared its performance to the regression-seq-dist model, which achieved the best performance on the regression task. Moreover, by creating the regression-dist model, we tested whether a machine-learning model can approximate a non-trivial function over the distance feature and generate predictions that result in improved average Pearson correlation compared to the Pearson correlation between the experimental off-target cleavage read counts and the distance feature, which we term *reads-distance-corr*.

The results on both CHANGE-seq and GUIDE-seq show that Pearson correlation achieved by the models trained only using the distance feature is higher than *reads-distance-corr*. First, we compared the performance of predicting the off-target cleavage efficiency on the CHANGE-seq dataset (Figure 5a). The regression-dist model achieved an average Pearson correlation of 0.439, which is higher than the reads-distance-corr with an average Pearson correlation of 0.389 (p-value= 7.29 · 10^-4^), and inferior to the regression-seq-dist model, which achieved 0.509 (p-value= 7.77 · 10^-8^). Second, we compared the performance of predicting the cleavage efficiency on the GUIDE-seq dataset (Figure 5b). The regression-dist model achieved an average Pearson correlation of 0.774, which is higher than the reads-distance-corr with an average Pearson correlation of 0.750 (p-value= 6.44 · 10^-5^), and on par with the regression-seq-dist model, which achieved 0.752 (p-value= 0.285). The full results, on each sgRNA, are reported in Supplementary Tables S1 and S4.

**Figure 5:**
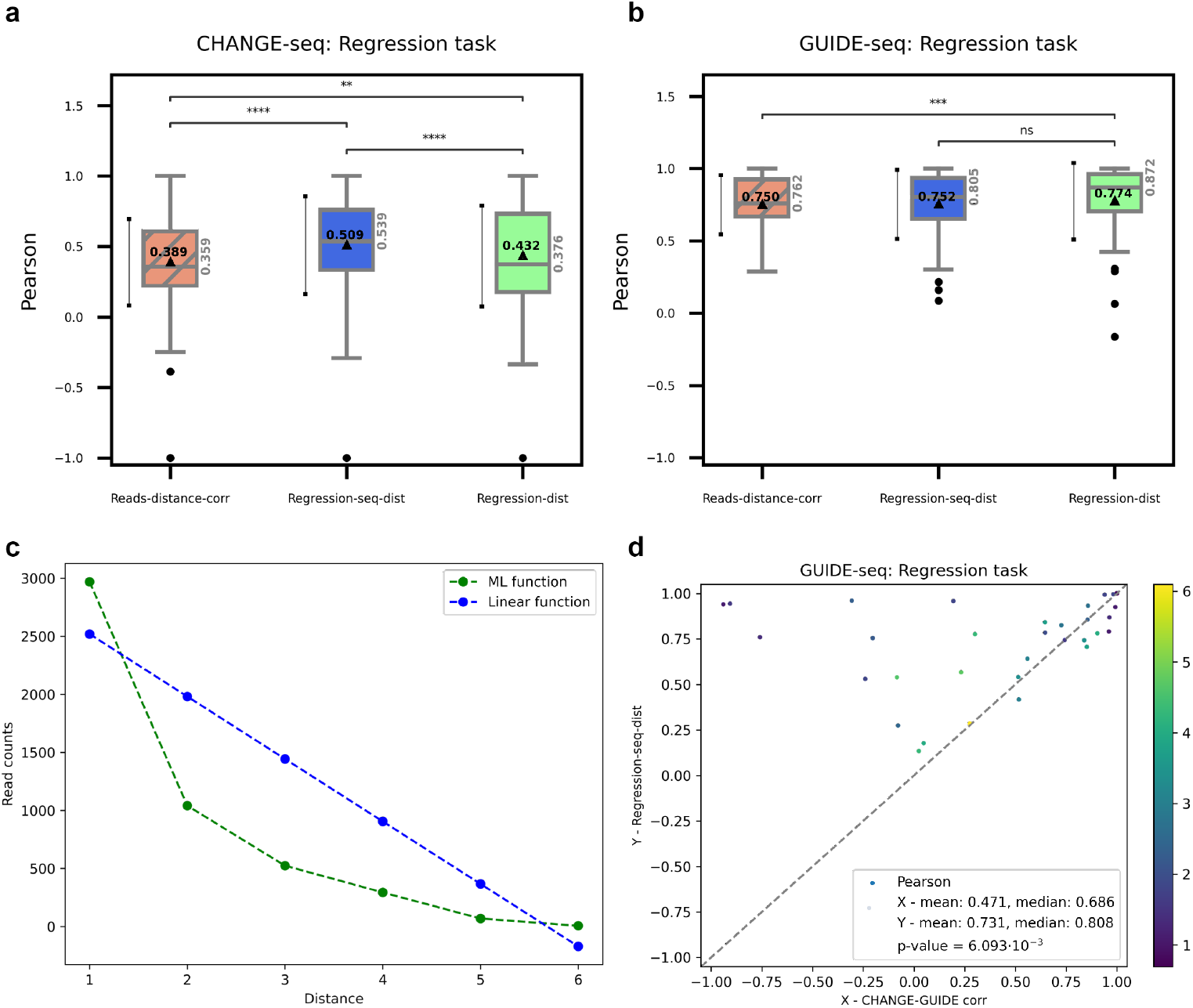
Machine-learning performance compared to experimental measurements. **(a-b)** Comparison of regression-seq-dist and regression-dist models in the regression task with the experimental reads-distance-corr. Performance was gauged by Pearson correlation of measured and predicted read counts. The average Pearson correlation is denoted by a triangle within each box plot. Error bar of one standard deviation is located on the left of each box plot. The median Pearson correlation value is reported in grey on the right of each box plot. Statistical significance via Wilcoxon signed-rank test is denoted by: ns 5o 10^-2^ < p ≤ 1; ** 1 · 10^-3^ < p ≤ 1 · 10^-2^; *** 1 · 10^-4^ < p ≤ 1 · 10^-3^; **** p ≤ 1 · 10^-4^. **(a)** Evaluation on the CHANGE-seq dataset. **(b)** Evaluation on the GUIDE-seq dataset. **(c)** The function of read counts vs. the distance learned by the regression-dist model compared to a linear fit of cleavage efficiency to the distance. **(d)** Comparison of prediction performance by CHANGE-seq measurements and the regression-seq-dist model on GUIDE-seq data. For this evaluation, the active off-target sites were considered as the active off targets that appear in both the CHANGE-seq and GUIDE-seq dataset. In same manner, the inactive set was determined. Using a leave-11-sgRNAs-out cross-validation, we evaluated the performance on each sgRNA individually. For each sgRNA, the Pearson correlation between the the CHANGE-seq and GUIDE-seq read counts (X-axis) is compared to the performance of the regression-seq-dist model in the regression task (Y-axis). Each point in the scatter plots represents performance evaluated on a sgRNA. The color of each point in the scatter plots represents the log of the number of active off-target experimentally detected for this sgRNA. The reported p-value was computed by Wilcoxon signed-rank test.

To examine the function, learned by the regression-dist model, of off-target cleavage efficiency vs. distance, we assembled the predictions of the 10 models trained in the leave-11-sgRNAs-out cross-validation by averaging the predictions for different distances. As comparison, we fitted for each sgRNA a linear regression, and averaged the predictions of the models for different distances. Interestingly, the regression-dist model was able to learn a more complex function than the linear fit (Figure 5c). We hypothesize that this function models the biophysical properties of RNA-DNA base-pairing in the Cas9-editing site.

### Evaluation of transfer learning from CHANGE-seq to GUIDE-seq

To further utilize the CHANGE-seq dataset for GUIDE-seq predictions, we propose, for the first time as far as we are aware of, the use of transferlearning techniques. We compared the performance of models based on transfer learning, one by adding trees, denoted GS-TL-add, and one by updating trees, denoted GS-TL-update, with a model trained only on the GUIDE-seq data, denoted GS, in both classification and regression tasks. For the evaluation of the classification task, we trained models equivalent to classification-seq and classification-seq-dist models. For the evaluation of the regression task, we trained models equivalent to regression-seq and regression-seq-dist models.

The models based on transfer learning outperformed the models trained only on GUIDE-seq data in classifying active off-target sites (Figures 6a,b). When training on all the 48 GUIDE-seq experiment, GS-TL-add achieved the best average AUPR of 0.573 for the model trained only with the sequence features, and 0.604 for the model trained with both the sequence and distance features. In the same test, GS-TL-update and GS models trained only with the sequence features achieved an average AUPR of 0.537 and 0.518, respectively. GS-TL-update and GS models trained with both the sequence and distance features achieved an average AUPR of 0.545 and 0.469, respectively. In addition, as expected from our previous analysis of models trained on the CHANGE-seq dataset, GS-TL-add model already achieves a high average AUPR even when the transfer learning is performed on only one GUIDE-seq experiment. Interestingly, GS-TL-update model achieves a low average AUPR with only one GUIDE-seq experiment; however, when adding more GUIDE-seq experiments to the training set, the performance improvement is almost instantaneous compared to GS model.

**Figure 6:**
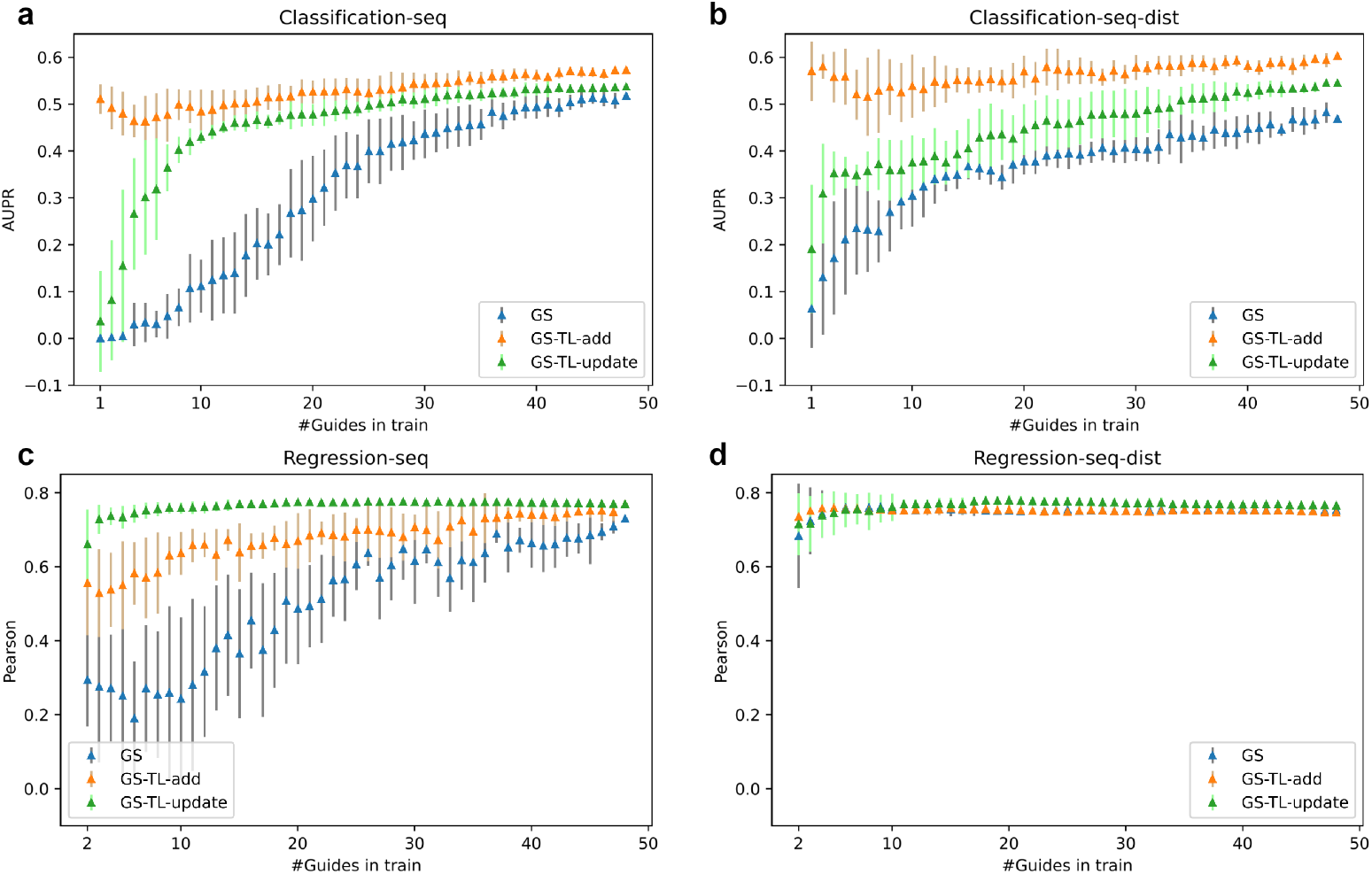
Transfer learning from CHANGE-seq to GUIDE-seq. **(a-d)** Comparison of GS (GUIDE-seq only), GS-TL-add (transfer learning by adding trees), and GS-TL-update (transfer learning by updating trees) models in both the classification and regression tasks. In each comparison, the effect of adding more GUIDE-seq experiments to the transfer-learning set was evaluated. The average performance and standard deviation over 10 evaluations, where in each evaluation the order of adding the experiments to the training set is different, is denoted by a triangle and vertical line, respectively. **(a-b)** Classification task performance comparison of GS, GS-TL-add, and GS-TL-update classification models trained with only sequence features (a), and with both the sequence and distance features (b). **(a-b)** Regression task performance comparison of GS, GS-TL-add, and GS-TL-update regression models trained with only sequence features (c), and with both the sequence and distance features (d).

When training on all 48 GUIDE-seq experiments, all models achieved similar performance in predicting off-target cleavage efficiency (Figures 6c,d). GS-TL-update achieved the highest average Pearson correlation of 0.770 for the model trained only with the sequence features and 0.766 for the model trained with both the sequence and distance features. In the same test, GS-TL-add and GS models trained only with the sequence features achieved an average Pearson correlation of 0.770 and 0.730, respectively. GS-TL-add and GS models trained with both the sequence and distance features achieved an average Pearson correlation of 0.747 and 0.753, respectively. In addition, in this task, adding more GUIDE-seq experiments to the training data for transfer learning had almost no effect on the performance. In particular, and in contrast the classification task, GS-TL-update has almost the same performance when transfer learning with only two GUIDE-seq experiments or with all 48 experiments. Moreover, when adding the distance feature to the features set, only a few GUIDE-seq experiments are required to train the GS model to achieve its best performance.

## Discussion

In this study, we utilized datasets with unprecedented scale and accuracy produced for both *in vitro* and *in vivo* off-target sites to perform a systematic evaluation of data processing and formulation of the CRISPR off-target site prediction problem. To achieve this goal, we evaluated various models to predict off-target sites and to predict off-target cleavage efficiency. We examined the models in different testing scenarios, and as a result, concluded fundamental insights in developing and training models for CRISPR off-target site prediction.

First, we looked for the optimal approach to train a regression model to predict cleavage efficiency. We found that a log transformation of the read counts, which reduced the skewness of data, is critical for improving prediction performance in both regression and classification tasks. In addition, we found that training regression models with inactive off-target sites improves the performance of the predictive models in the regression task when evaluated only on the active off-target set. As expected, it also improves the model’s ability to perform prediction for the classification task.

Second, we tested what are the best predictive models for the classification and regression tasks when trained and tested on an *in vitro* dataset (i.e., CHANGE-seq). In addition, we tested how the predictive models generalize to an *in vivo* dataset (i.e., GUIDE-seq). We found that the use of a binary classifier trained with sequence and distance features achieved the best classification performance in both *in vitro* and *in vivo* datasets. Surprisingly, we observed that a regression model trained with sequence and distance features can successfully solve the classification task, and it achieved performance on par with a classification model on the *in vivo* dataset. In the regression task, we found that a regression model trained with sequence and distance features resulted in the best performance on both datasets. Interestingly, we noticed that a classifier trained only with sequence features predicted more accurately than an equivalent model trained with both the sequence and the distance features. We hypothesize that adding the distance feature to the classifier enhances the ability of the model to make a hard decision in the classification task, but can hamper its ability to predict off-target cleavage efficiencies, which is analog to a soft decision.Overall, we concluded that adding the distance feature improves the prediction for both tasks. Strikingly, we showed that the predictive models trained on an *in vitro* dataset could be generalized to an *in vivo* dataset, and we provided a proof-of-concept that our machinelearning approach is superior to the experimental relation between the two datasets (Figure 5d and Supplementary Figure S3).

Third, we studied the impact of the distance feature by training off-target cleavage efficiency regression models with only the distance feature. We observed that the machine-learning models achieved a higher Pearson correlation in both datasets than the Pearson correlation between the experimental off-target cleavage read counts and the distance feature. More importantly, we discovered that when testing the generalization of the regression models on the *in vivo* dataset, the regression model trained only with the distance feature obtained results on par with the regression model trained with both the sequence and the distance features. This might indicate that when trying to generalize off-target cleavage efficiency prediction from *in vitro* to *in vivo*, the most important feature is the distance.

Fourth, to overcome the problem of training a model on an *in vivo* dataset, which usually contains a small number of active off-target sites, we utilized the *in vitro* CHANGE-seq dataset to train a model for the *in vivo* GUIDE-seq using transfer-learning techniques. Our models based on transfer-learning were superior to the models trained only on the GUIDE-seq dataset or only on the CHANGE-seq dataset. In addition, we showed that using transferlearning techniques, and in particular for the classification task, the amount of *in vivo* experiments needed for transfer learning is smaller than training without any transfer learning. Moreover, we observed that in the regression task, only a few *in vivo* experiments are needed, especially when we added the distance as an input feature to the model.

Several aspects of our study require further research in the future. First, it will be interesting to repeat the evaluations performed in this study while considering off-targets with DNA/RNA bulges. Following such evaluation, one can develop a computational model to predict off-target sites with improved performance compared to the state of the art [30]. Second, the efficiency of assays for measuring the genome-wide activity of CRISPR/Cas9 nucleases *in vivo* is limited by chromatin accessibility and other epigenetic factors [36]. Therefore, including such information in our evaluation may provide more insights regarding their addition to computational models for future improvement. Third, the datasets containing active and inactive off-targets sites are extremely imbalanced. In this study, we applied one solution to the imbalance problem. It will be interesting to test various solutions to training on imbalanced data in future research to find the optimal one.

## Conclusion

To conclude, we performed an extensive investigation of approaches to train and evaluate models for predicting off-target sites and off-target cleavage efficiencies. We demonstrated that machine-learning models are capable of utilizing large-scale datasets to learn key features of off-target sites. We showed that our predictive models trained on *in vitro* data are able to make predictions on *in vivo* data more accurately then the *in vitro* measurements themselves. We are confident that any study that will aspire to develop a new off-target predictor will find our insights invaluable.

## Data Availability

The software and code are publicly available via github.com/OrensteinLab/SysEvalOffTarget.

## Supplemental Data

### Supplementary Figures

Supplementary Figures S1-3.

### Supplementary Tables S1-S4

Prediction performance on each CHANGE/GUIDE-seq sgRNA of the various models developed based on CHANGE-seq dataset.

## Acknowledgements

We would like to thank our colleagues at the CRISPR-IL consortium for thoughtful discussions over the off-target prediction problem.

## Author contributions

O.Y. conducted all the analyses and developed all the code used through this study. M.A. generated preliminary results for this study. Y.O. supervised this study. All authors contributed to the writing of the manuscript.

