## Supplementary Figures for "A systematic evaluation of data processing and problem formulation of CRISPR off-target site prediction"

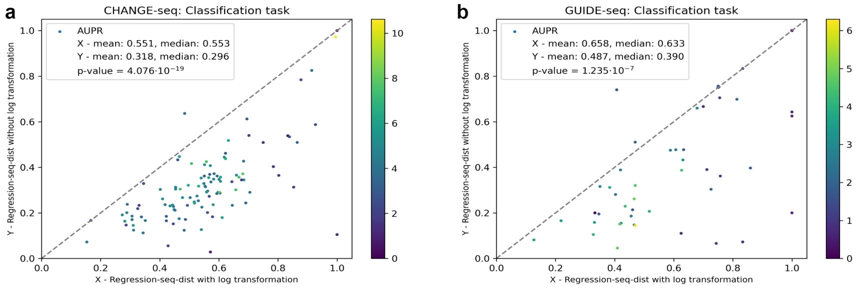


**Supplementary Figure S1: Read counts log transformation evaluation. (a-b)** Comparison of the regression-seq-dist model trained with log transformation and without. Using a leave-11-sgRNAs-out cross-validation, we evaluated the performance for the regression task on each sgRNA individually. Performance evaluation is preformed on the CHANGE-seq dataset (a), and the GUIDE-seq dataset (b). Performance was gauged by AUPR. Each point in the scatter plots represents performance evaluated on a sgRNA. The color of each point in the scatter plots represents the log of the number of active off-target sites experimentally detected for this sgRNA. The reported p-value was computed by Wilcoxon signed-rank test.

**Supplementary Figure S2: Training on both inactive and active off-target sites evaluation. (a-b)** Comparison of the regression-seq-dist model trained only on active off-target sites and trained on both active and inactive off-target sites. Using a leave-11-sgRNAs-out cross-validation, we evaluated the performance for the regression task on each sgRNA individually. Performance evaluation is preformed on the CHANGE-seq dataset (a), and the GUIDE-seq dataset (b). Performance was gauged by AUPR. Each point in the scatter plots represents performance evaluated on a sgRNA. The color of each point in the scatter plots represents the log of the number of active off-target experimentally detected for this sgRNA. The reported p-value was computed by Wilcoxon signed-rank test.

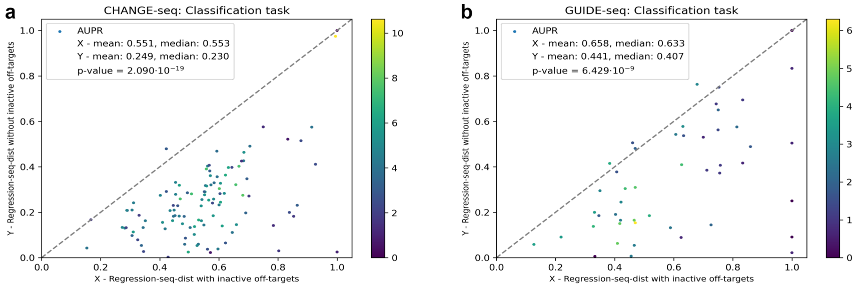

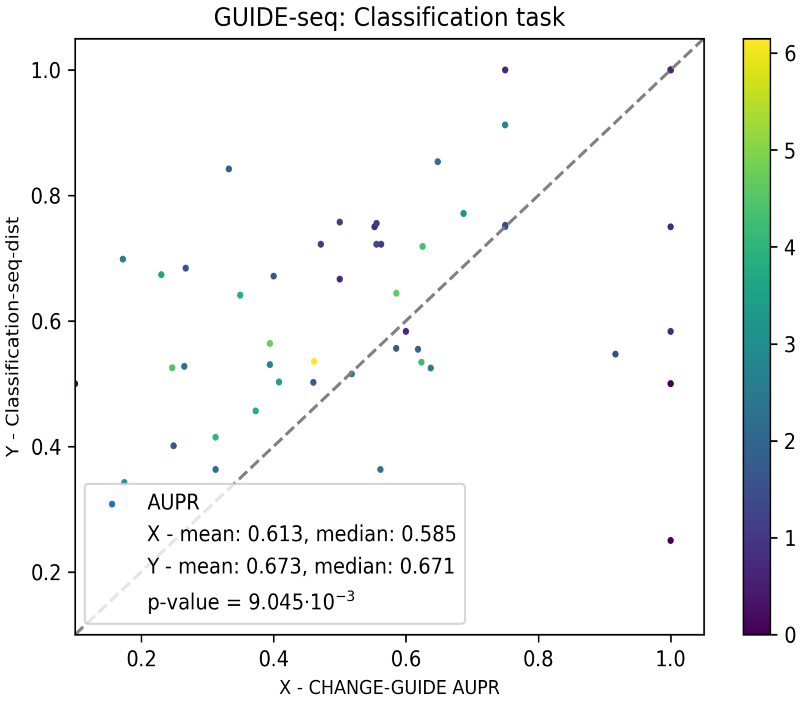


**Supplementary Figure S3: Machine-learning performance compared to experimental measurements.** Comparison of prediction performance by CHANGE-seq measurements and the classification-seq-dist model on GUIDE-seq data. For this evaluation, the active off-target sites were considered as the active off targets that appear in both the CHANGE-seq and GUIDE-seq dataset. In same manner, the non-active set was determined. Using a leave-11-sgRNAs-out cross-validation, we evaluated the performance on each sgRNA individually. For each sgRNA, the computed AUPR of the CHANGE-seq read counts and GUIDE-seq classification labels (X-axis) are compared to the performance of the classification-seq-dist model in the classification task (Y-axis). Each point in the scatter plots represents performance evaluated on one sgRNA. The color of each point in the scatter plots represents the log of the number of active off-target experimentally detected for this sgRNA. The reported p-value was computed by Wilcoxon signed-rank test.
